# Pollen contamination and mating structure in maritime pine (*Pinus pinaster* Ait.) clonal seed orchards revealed by SNP markers

**DOI:** 10.1101/2022.09.27.509769

**Authors:** Laurent Bouffier, Sandrine Debille, Pierre Alazard, Annie Raffin, Patrick Pastuszka, Jean-François Trontin

## Abstract

Maritime pine (*Pinus pinaster* Ait.) is a major forest tree species in south-western Europe. In France, an advanced breeding program for this conifer species has been underway since the early 1960s. Open-pollinated seed orchards currently supply more than 90% of maritime pine seedlings for plantation forestry. However, pollen contamination and mating structure have been poorly studied in such seed orchards whereas they could impact genetic gains and diversity. We analyzed these features in three maritime pine clonal seed orchards. We addressed biological (tree genotype, age, flowering phenology) and environmental factors (vicinity with external pollen sources, orchard structure, soil type, climatic conditions) that are expected to determine the genetic composition of seed lots. Genetic analyses were based on an optimized set of 60 SNP markers and performed on 2,552 seedlings with Cervus software (likelihood inference methodology).

Pollen contamination rates were highly variable between seed lots (from 20 to 96%), with a mean value of 50%. Interpretative factors included the distance between the seed orchard and external pollen sources, rain during the pollination period, seed orchard age, soil conditions and seed parent identity. All parental genotypes from the seed orchards contributed to the offspring as pollen parents, but differences in paternal reproductive success were detected. Finally, the overall self-fertilization rate was estimated at 5.4%, with considerable variability between genotypes (from 0% to 26%). These findings are useful to formulate recommendations for seed orchard management (seed orchard location, soil and climate optimal conditions, minimum age for commercial seed lots harvesting) and for identifying new research perspectives (exploring links between pollen contamination and climatic data, genetic control of flowering traits).

## Introduction

The first breeding programs for forest trees were initiated in the mid-20^th^ century, to improve forest genetic resources in terms of productivity, wood quality and adaptation to environmental conditions (Burdon et al. 2008; McKeand et al. 2003; Pâques 2013). These programs have been shaped by environmental, biological, economic, institutional and sociopolitical factors (Mullin and Lee 2013). However, they have always been constructed around two main populations (Namkoong et al. 1988): a breeding population initiated by selecting superior trees, generally managed over successive cycles of crossing-testing-selection (recurrent selection scheme), and a deployment population (improved Forest Reproductive Material - FRM -) released for commercial plantations. The breeding population carries a high level of genetic diversity, to limit inbreeding (targeted effective size of 30 to 70) and ensure future genetic gains (Danusevicius and Lindgren 2005). By contrast, the deployment population is selected to maximize genetic gains for selection criteria. The genetic diversity of deployment populations varies between breeding programs, from single genotypes (clonal forestry) or mixtures of selected genotypes (multiclonal forestry, also known as multivarietal forestry, Weng et al. 2011) to synthetic populations produced through sexual reproduction in open-pollinated seed orchards. The absence of contaminating pollen from surrounding stands is a necessary condition to achieve the expected genetic gains in such synthetic population. However, many studies, based on pedigree reconstruction, have suggested that there may be significant levels of pollen contamination in conifer seed orchards: they were carried out initially with allozymes (Harju and Nikkanen 1996; Yazdani and Lindgren 1991) or RAPD (Goto et al. 2002), and subsequently with microsatellites (Slavov et al. 2005; Torimaru et al. 2009). The recent development of single-nucleotide polymorphisms (SNP) has opened up new opportunities for analyzing more precisely mating patterns in seed orchards. In this article, we estimated pollen contamination and parental contributions in maritime pine seed orchards using an SNP array previously optimized by Vidal et al. (2015) for pedigree reconstruction.

Maritime pine (*Pinus pinaster* Ait.), which covers 4.2 million hectares, is a major forest tree species in south-western Europe (Abad Viñas et al. 2016). It accounts for 7% of the forested area in France (1.03 million ha), mostly in the large Landes de Gascogne forest (0.81 million ha), but it provides 18% of lumber and 27% of pulpwood production nationally (FCBA 2020). A breeding program was initiated in the 1960s from a base population selected in South-West France for two main criteria: growth rate and stem straightness. The breeding population was subjected to three cycles of recurrent selection (Mullin and Lee 2013). Over the past ten years (2011-2021), 375 million improved seedlings were sold by forest nurseries (French Ministry of Agriculture 2022), making maritime pine the leading species for plantation forestry in France. The seedlings were obtained from seeds collected in open-pollinated clonal and polycross seed orchards (Baradat 1987) displaying an expected genetic gain of 30% for breeding objective traits. However two studies (Plomion et al. 2001; Plomion et al. 2005) have reported high level of pollen contamination in maritime pine seed orchards which could scale down this gain. A better understanding of the reproductive regime operating in these seed production structures is therefore essential to improve management practices for maximizing genetic gains.

We hypothesized that the orchard’s proximity with external pollen sources constituted by the plantations from the maritime pine Landes de Gascogne forest was a key explanatory factor for pollen contamination. However many other environmental and biological factors can be highlighted such as: the orchard age, the orchard soil type, the climatic conditions of the pollination year (especially rainfall during the pollination period), the orchard structure (number of ramets per parental genotype and their localization within the orchard), the flowering phenology of the parental genotypes. We considered various sampling strategies (2,552 seedlings in total) based on three maritime pine open-pollinated clonal seed orchard (CSO) to explore the role of these explanatory factors for pollen contamination, parental contributions (including self-fertilization) and genetic diversity. We discuss the results from the perspective of optimizing the deployment of new seed orchards of this key tree species for plantation forestry in France.

## Materials and methods

### Plant material

Sampling was carried out in third-generation CSO currently producing maritime pine seed lots (improved FRM) by open pollination. For each seed collected, the seed parent (maternal genotype) is known, as the parental genotypes are identified in the field, but the pollen parent (paternal genotype) is unknown, due to the system of open pollination. We studied seeds from three CSO established between 2002 and 2006. Each CSO was composed of the same 50 selected genotypes deployed through grafting. An additional genotype has been introduced following a restocking operation after establishment of the seed orchards. The actual contribution of each genotype varies within and between CSO due to differences in the number of ramets per genotype. The three CSO (Table 1) differ principally in terms of their location and soil characteristics:

**Table 1.** Characteristics of the three maritime pine clonal seed orchards (CSO) sampled.

| **Code** | **Name**<br>**(reference)** | **Establishment**<br>**year** | **Nearest**<br>**maritime**<br>**pine stands** | **Soil type** | **Area**<br>**(ha)** | **No. genotypes**<br>**(no. trees)** |
| --- | --- | --- | --- | --- | --- | --- |
| CSO-1 | Saint-Laurent2-VF3<br>(PPA-VG-014) | 2006 | < 500 m | Sandy | 15.0 | 46<br>(3171) |
| CSO-2 | Beychac-VF3<br>(PPA-VG-011) | 2002-2003 | ~ 5 km | Clay loam | 15.5 | 47<br>(3676) |
| CSO-3 | Saint-Sardos VF3<br>(PPA-VG-015) | 2003 | > 20 km | Clay loam | 6.5 | 48<br>(1565) |

– CSO-1, established in the northern part of the Landes de Gascogne forest (i.e. surrounded by maritime pine plantations) on sandy soils;

– CSO-2, established on the eastern outskirts of the Landes de Gascogne forest (i.e. nearest maritime pine plantations a few kilometers away) on clay loam;

– CSO-3, established at the southernmost eastern location, outside the Landes de Gascogne forest (i.e. nearest maritime pine plantations more than 20 kilometers away) on clay loam.

In total, 2,552 seedlings were considered, through three sampling strategies (Table 2):

**Table 2.** Sampling strategies (SS1, SS2 and SS3) of 2,552 seedlings in three maritime pine clonal seed orchards (CSO) over three pollination years. Seed orchard age is calculated from the time of the CSO plantation (which occurs one year after the grafting of the scion on a two years old rootstock).

| Seed orchard | Pollination year | Seed orchard age (yr.) | Sampling strategy | Sampling zone | No. seed parent genotypes* | No. seedlings genotyped (no. seedlings per genotype) |
| --- | --- | --- | --- | --- | --- | --- |
| CSO-1 | 2011 | 5 | SS1 | Center | 4 (A, B, C, D) | 240 (60) |
|  |  |  |  | Border | 4 (A, B, C, D) | 240 (60) |
|  | 2013 | 7 | SS1 | Center | 4 (A, B, C, D) | 116 (27-30) |
|  |  |  |  | Border | 4 (A, B, C, D) | 120 (30) |
|  | 2014 | 8 | SS3 | Unknown | Unknown | 147 (unknown) |
| CSO-2 | 2011 | 8-9 | SS1 | Center | 4 (A, B, C, D) | 240 (60) |
|  | 2013 | 10-11 | SS1 | Center | 4 (A, B, C, D) | 238 (58-60) |
|  | 2013 |  | SS2 | Center | 20 | 590 (27-30) |
|  | 2014 | 11-12 | SS3 | Unknown | Unknown | 142 (unknown) |
| CSO-3 | 2011 | 8 | SS1 | Center | 4 (A, B, C, D) | 240 (60) |
|  | 2013 | 10 | SS1 | Center | 4 (A, B, C, D) | 90 (17-29) |
|  | 2014 | 11 | SS3 | Unknown | Unknown | 149 (unknown) |

\* A, B, C, D: reference genotypes with early (A, B) or late (C, D) female flowering (4-6 cones collected each year on the same two ramets per genotype and sampling zone).

– Sampling strategy 1 (SS1): in the fall of 2012 and 2014 (pollination years 2011 and 2013), one central sampling zone was defined per CSO, except for CSO-1, in which two zones were considered (center vs. border). Four genotypes (denoted by A, B, C and D) were selected according to seed parent flowering phenology (Trontin et al. 2019): two early (A, B) and two late flowering genotypes (C and D). Each year, four to six cones were collected from two ramets (the same ramets were sampled in 2012 and 2014) per genotype and per sampling zone. After germination, 17 to 60 seedlings per genotype (1,524 in total) were sampled at random.

– Sampling strategy 2 (SS2): in the fall of 2014 (pollination year 2013), three cones were collected for 20 additional genotypes, randomly selected from CSO-2, and 27 to 30 seedlings per genotype were sampled at random (590 in total).

– Sampling strategy 3 (SS3): commercial seed lots, i.e. seeds extracted from bulked cones collected from 40 randomly selected trees from each CSO, were sampled in the fall of 2015 (pollination year 2014) and 142 to 149 seedlings per CSO were sampled (438 in total).

For each lot harvested, the seeds were germinated and grown in greenhouse conditions for 6 months. Seed parent identity was recorded for each seedling in SS1 and SS2, whereas the identities of both seed and pollen parents were unknown in SS3. There is a one-year time lag between pollination and fertilization in maritime pine. To avoid confusion, the years specified for each seed lots hereafter are the pollination years and not the sampling (fertilization) years.

### DNA extraction and genotyping

Needle tissues from the 2,552 six-month-old seedlings described above and from the 51 seed orchard parental genotypes (two ramets sampled per genotype) were ground to a fine powder in liquid nitrogen and subjected to DNA extraction with the Qiagen DNeasy® 96 Plant Kit, in accordance with the manufacturer’s protocol. The DNA was quantified with a NanoDrop microvolume spectrophotometer (Thermo Fisher Scientific Inc., Waltham, CA, USA) and diluted to 10 ng/μl. DNA samples were then genotyped with the 80 SNP markers developed by Vidal et al. (2015). Genotyping was performed with the Sequenom MassARRAY iPLEX Gold assay (Sequenom, San Diego, CA, USA). SNP markers were discarded if genotype calling was unsuccessful for more than 10% of the samples or if they deviated from Hardy-Weinberg equilibrium.

### Pedigree reconstruction

Pedigree reconstruction was performed by likelihood inference with Cervus 3.0.7 (Kalinowski et al. 2007). Paternity reconstruction analysis was performed when the seed parent was known (SS1 and SS2). By contrast, parental reconstruction analysis was performed when both seed and pollen parents were unknown (SS3). We assumed a 0.1% genotyping error rate (estimation based on repeated genotyping of the 51 parental genotypes). The delta score (i.e. the difference in LOD scores of the two most likely candidate parents) was used as a criterion for the assignment of paternity with 99% confidence. The critical values of delta scores were determined from simulations of 100,000 offspring. We allowed only one mismatched allele between a given offspring and its parents. A seedling was considered to result from pollen contamination (pollination by a pollen grain originating from outside the seed orchard) if no pollen parent from the 51 parental genotypes was found in the paternity reconstruction analysis or if only one parent was identified in the parental reconstruction analysis.

### Parental contribution

The paternal contribution for a given genotype was estimated, for SS1 and SS2, as the number of seedlings in which the pollen parent was identified divided by the total number of pollen parents recovered. This estimate was compared to a theoretical paternal contribution to assess the deviation from equal paternal contributions. The theoretical paternal contribution for a given genotype *i* was calculated with weighting according to the number of ramets per CSO, as follows:

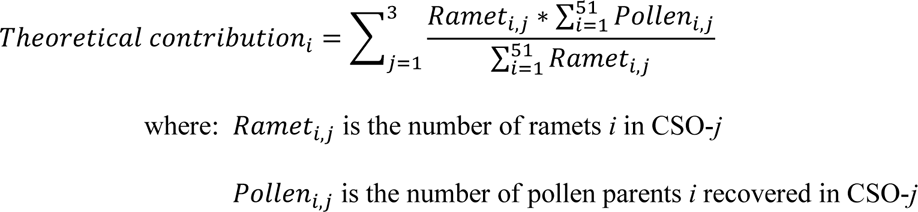

The self-fertilization rate was estimated by dividing the number of seedlings with two identical parental genotypes by the total number of seedlings for which both parental genotypes were recovered.

The significance of frequency differences for contamination rate and parental contribution was estimated with a chi-squared test of homogeneity (α = 0.05).

### Genetic diversity parameters

As described above, the seed parent genotypes were clonally represented, within the three CSO, by different numbers of ramets. The census number of seed parents (N) per CSO was, therefore, different from the effective number of seed parents (N_eff_) defined by 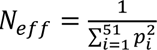 where p*i* is the contribution of genotype *i* (Kang et al. 2001). The effective number was also calculated for the contribution of the pollen parent, initially without considering pollen contamination (N_eff, pollen parent_) and then considering each immigrant pollen grain as a unique pollen parent (N_eff, pollen parent*_).

## Results

### Genotyping and pedigree reconstruction

We retained 60 of the 80 SNP tested for pedigree reconstruction based on the genotyping call restriction and Hardy-Weinberg equilibrium criteria. The mean polymorphic information content (Hearne et al. 1992) per marker was 0.372. The mean non-exclusion probability (Marshall et al. 1998), defined as the probability of not excluding a candidate parent of a given offspring, was 4 x 10^-4^, and the probability of genotypes not differing between two randomly chosen individuals was 4 x 10^-26^. The 2,552 seedlings were successfully genotyped for 35 to 60 SNP (mean of 56.8 SNP per seedling). After concatenation of the genotyping data obtained from two different ramets, data for 59 to 60 SNP were available for the 51 seed orchard parental genotypes.

All seed parent identities were confirmed for seeds collected in SS1 and SS2. In addition, based on paternity analyses, a pollen parent was identified from among the 51 parental genotypes for 1,023 of the 2,114 samples collected from a known seed parent (48.4%). These samples included 57 samples arising from the self-pollination of parental genotypes (5.6%). A parental analysis was performed on the commercial seed lots (SS3: 438 samples): both parents were identified from the 51 parental genotypes for 264 samples (60.3%), 12 of which were generated by self-fertilization (4.5%), and only one parent was recovered for the remaining 174 samples (39.7%).

### Pollen contamination

Overall, pollen contamination rates was estimated at 50% for the 2,552 samples analyzed: 558 pollen parents were recovered for 1524 samples in SS1, 465 pollen parents were recovered for 590 samples in SS2 and both seed and pollen parents were recovered for 264 of 438 samples in SS3. Pollen contamination rates are expressed by CSO and by pollination year in Figure 1a (SS1 and SS3). Whatever the pollination year considered, contamination rates were significantly higher in CSO-1 than in CSO-2 and CSO-3 (CSO-2 and CSO-3 differed significantly in 2013, but not in 2011 and 2014).

**Figure 1.**
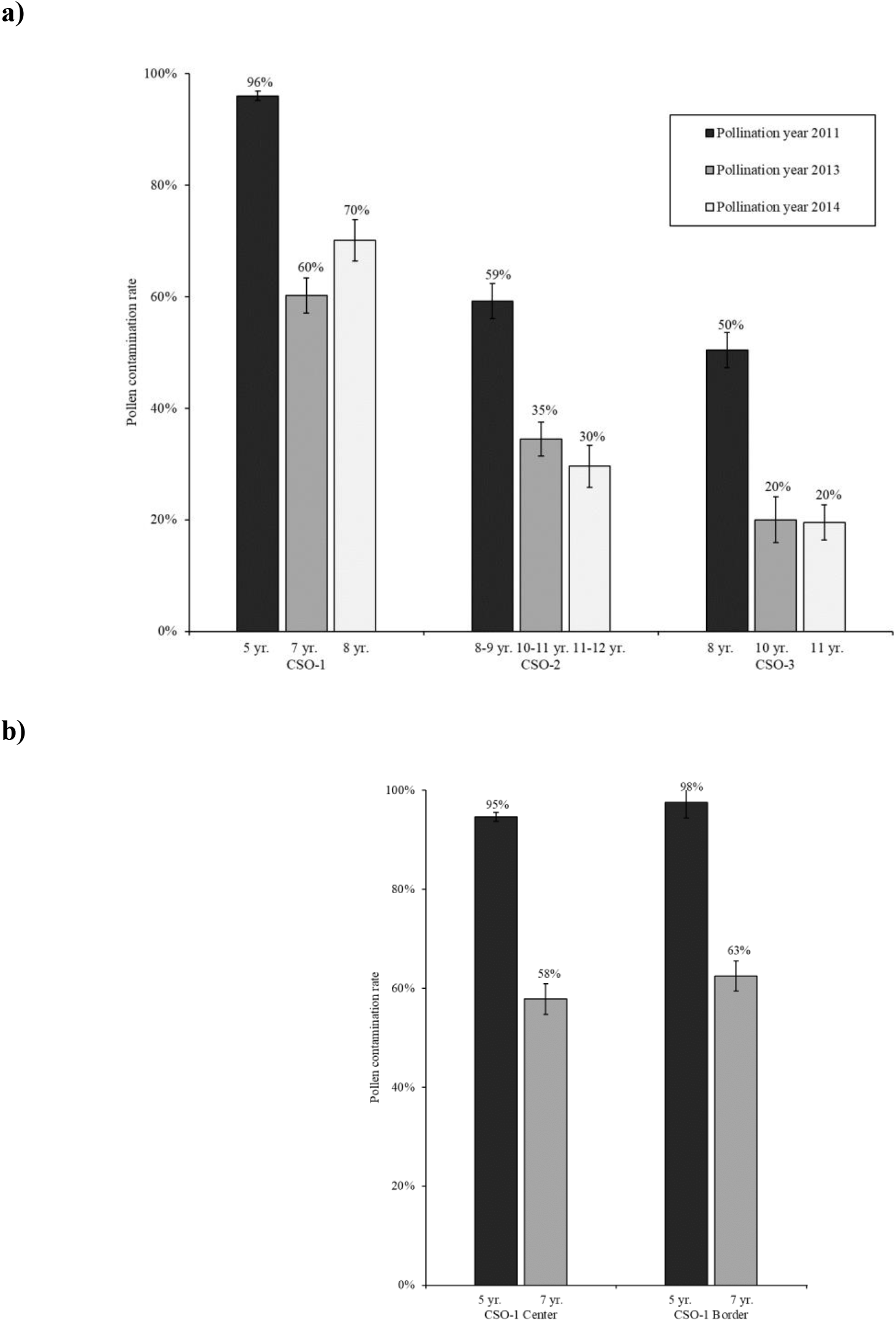
Pollen contamination rates (%) observed: **a)** in the three maritime pine clonal seed orchards (CSO-1, CSO-2, CSO-3) over three pollination years (2011, 2013, 2014), **b)** within the clonal seed orchard CSO-1 for two sampling zones (center, border) over two pollination years (2011 and 2013). Pollination years 2011 and 2013 corresponds to the sampling strategy SS1 (seeds collected on the four reference seed parents A, B, C, D). Pollination year 2014 corresponds to the sampling strategy SS3 (commercial seed lots from unknown seed parents) (see Table 2). The seed orchard age is reported for each pollination year. Bars: standard errors.

For a given CSO, contamination rates were significantly higher in 2011 than in 2013 and 2014 (no significant difference was found between 2013 and 2014). The pollen parent originated from outside the CSO-1 orchard for 96% of the samples collected in 2011 vs. 60% in 2013 and 70% in 2014. A similar inter-annual trend was observed in CSO-2 (59% vs. 35% and 30%, respectively) and CSO-3 (50% vs. 20% and 20%, respectively). It is important to note that the pollination year effect is partly confounded, in this study, with the CSO age (also reported for each CSO in Figure 1).Within CSO-1, two sampling zones were considered (central vs. border) but the pollen contamination rates were not significantly different for either 2011 or 2013 (Figure 1b).

No significant effect of seed parent identity on contamination rate was found when considering SS1 (the contamination rates estimated over the three CSO was 62.1%, 61.0%, 61.9% and 68.7% for seed parents A, B, C and D, respectively, see Table 3). However, when analyzing a higher number of seed parents (the 20 seed parents investigated in SS2 and for the four seed parents A, B, C, D in SS1) for the pollination year 2013 in CSO-2 (Figure 2), the variability of pollen contamination rates was high and depended on seed parent identity, ranging from 10% to 45% (the mean value over the 24 seed parents identities was 25%).

**Figure 2.**
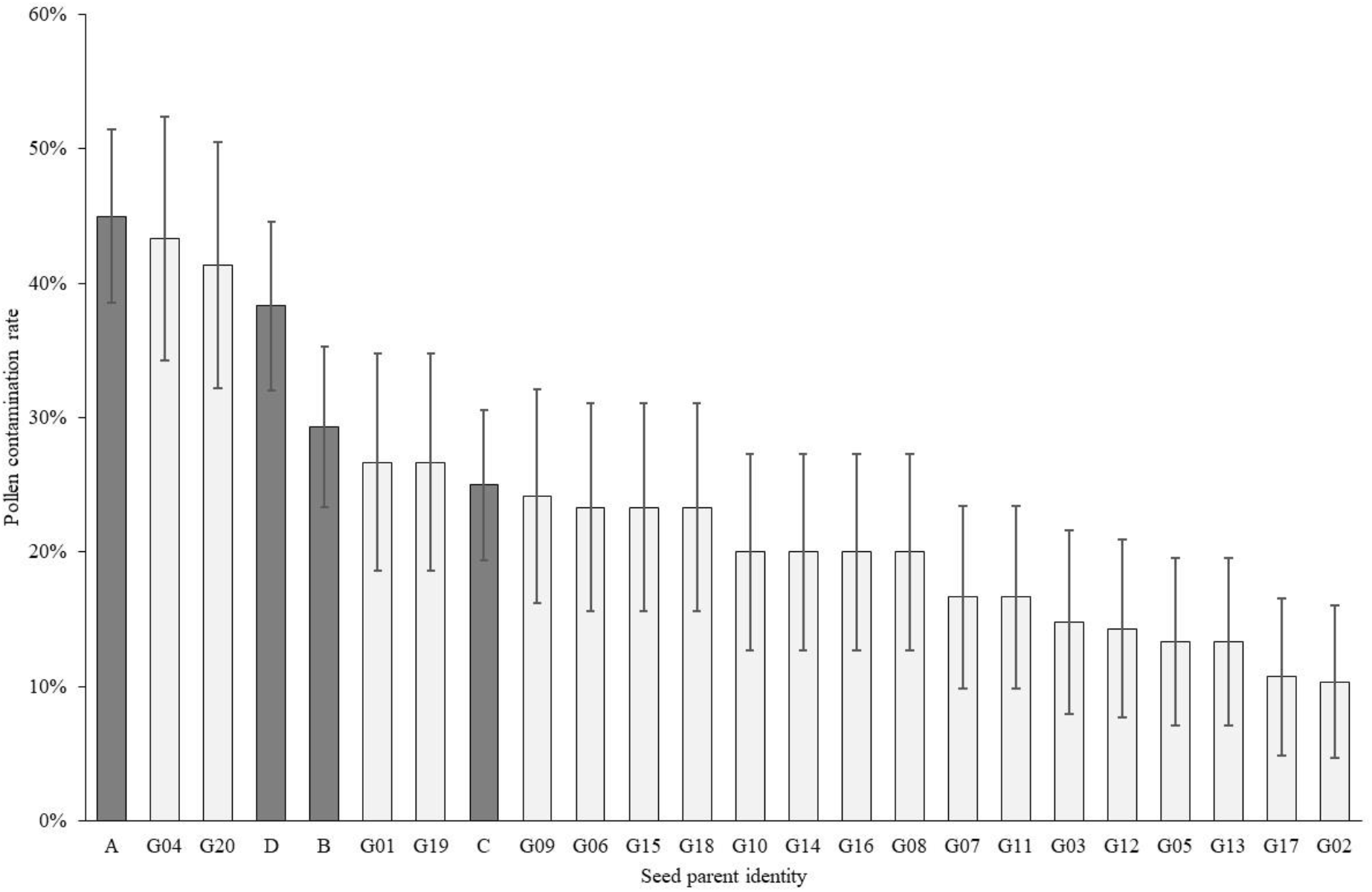
Pollen contamination rates (%) on 24 seed parent genotypes of maritime pine clonal seed orchard CSO-2 pollinated in 2013. Dark grey: four reference seed parents (sampling strategy SS1); Light grey: 20 seed parents (sampling strategy SS2); Bars: standard errors.

**Table 3.**
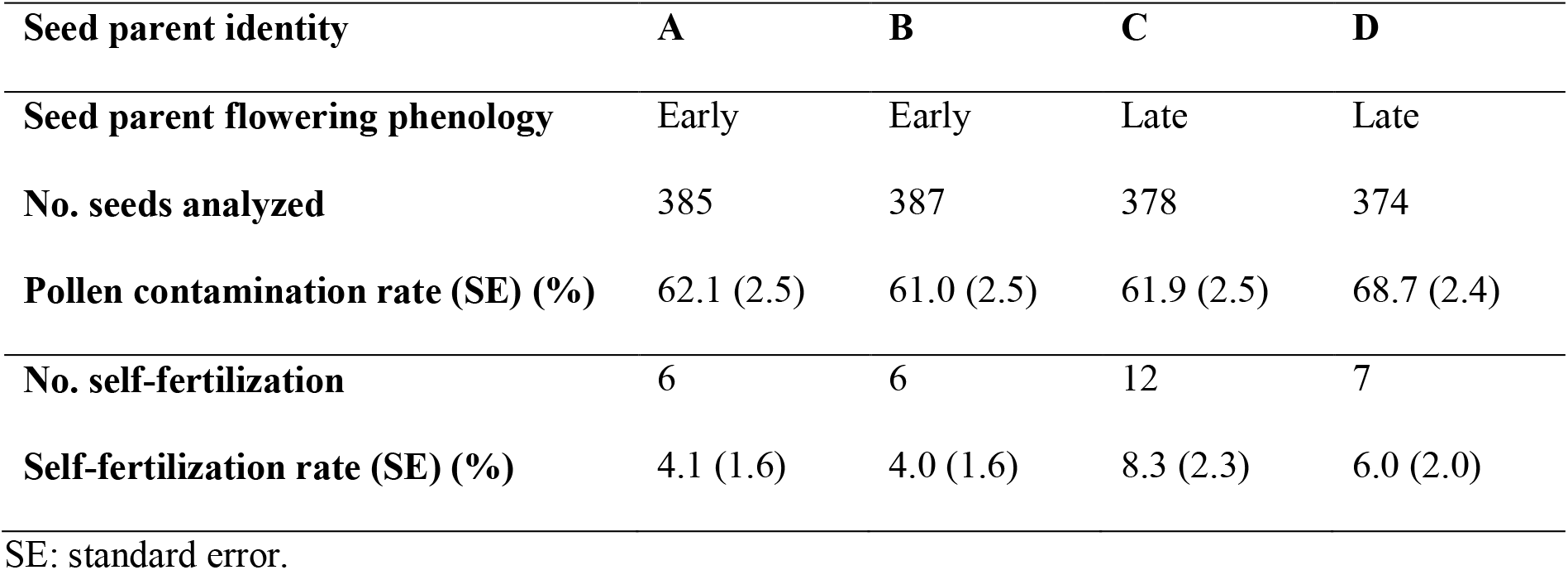
Pollen contamination and self-fertilization rates per reference seed parent genotype in three maritime pine clonal seeds orchards (CSO-1, CSO-2, CSO-3) over 2011 and 2013 (sampling strategy SS1, see Table 2).

### Paternal contribution

It was possible to estimate paternal contributions only with SS1 and SS2, for which the seed parent identity was known. These contributions are reported, by CSO, for each genotype in Figure 3. All parental genotypes contributed as pollen parents, but a high degree of heterogeneity was observed (the 51 parental genotypes were identified as pollen parents from 1 to 28 times). As SS2 focused exclusively on CSO-2, a larger number of pollen parents were recovered for CSO-2 (719) than for CSO-1 (113) and CSO-3 (191). The number of ramets per genotype and per CSO partly accounted for the heterogeneity of paternal contributions (Figure 4). Paternal contribution was, indeed, significantly correlated with genotype representativeness (expressed as the percentage of ramets per genotype in orchard); Pearson’s correlation coefficient was significant and estimated at 0.45 in CSO-1, 0.52 in CSO-2 and 0.48 in CSO-3.

**Figure 3.**
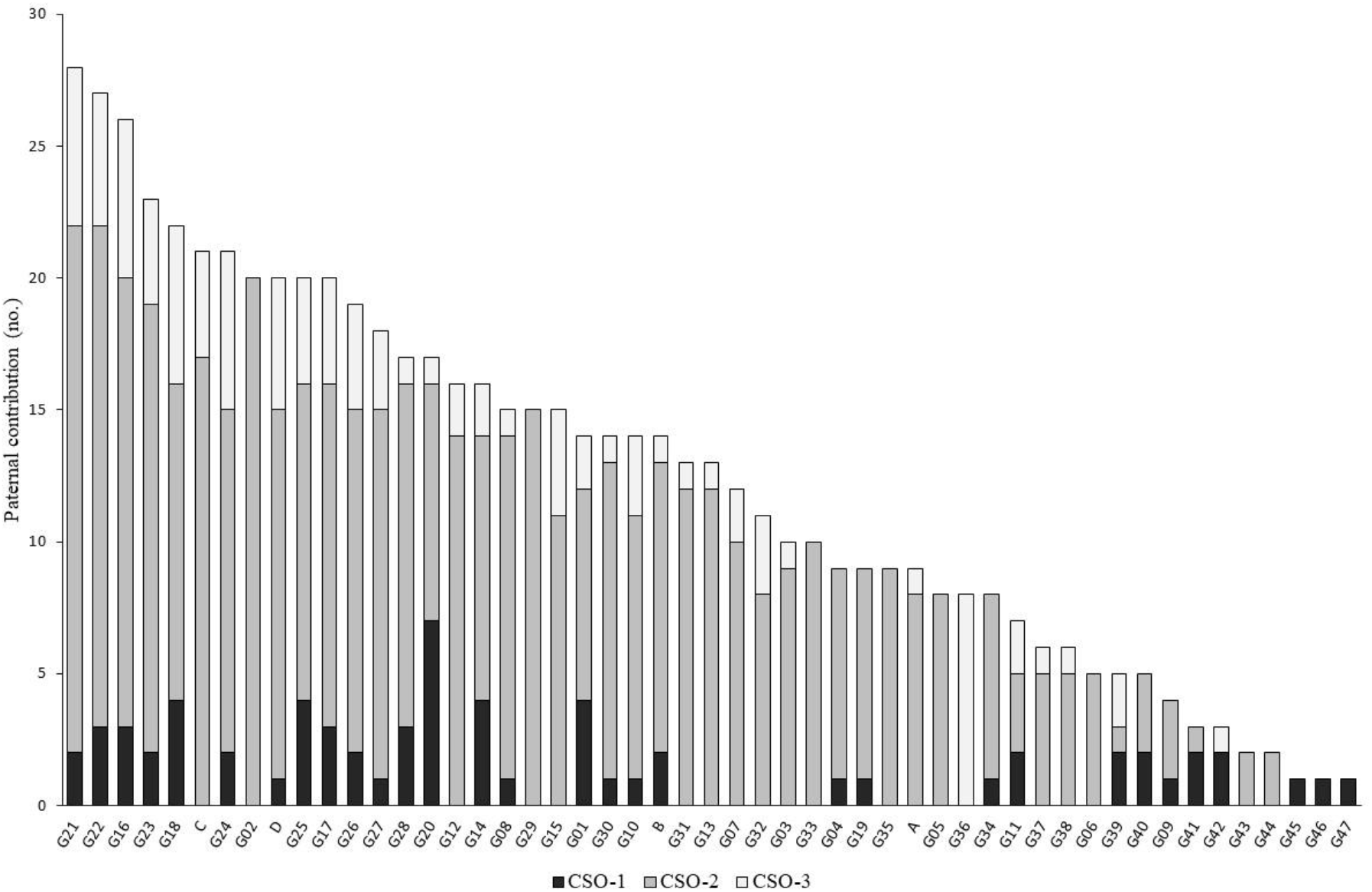
Paternal contribution (number of pollen parents) of each genotype observed in three maritime pine clonal seed orchards (CSO) over 2 pollination years (2011, 2013, sampling strategies SS1 and SS2).

**Figure 4.**
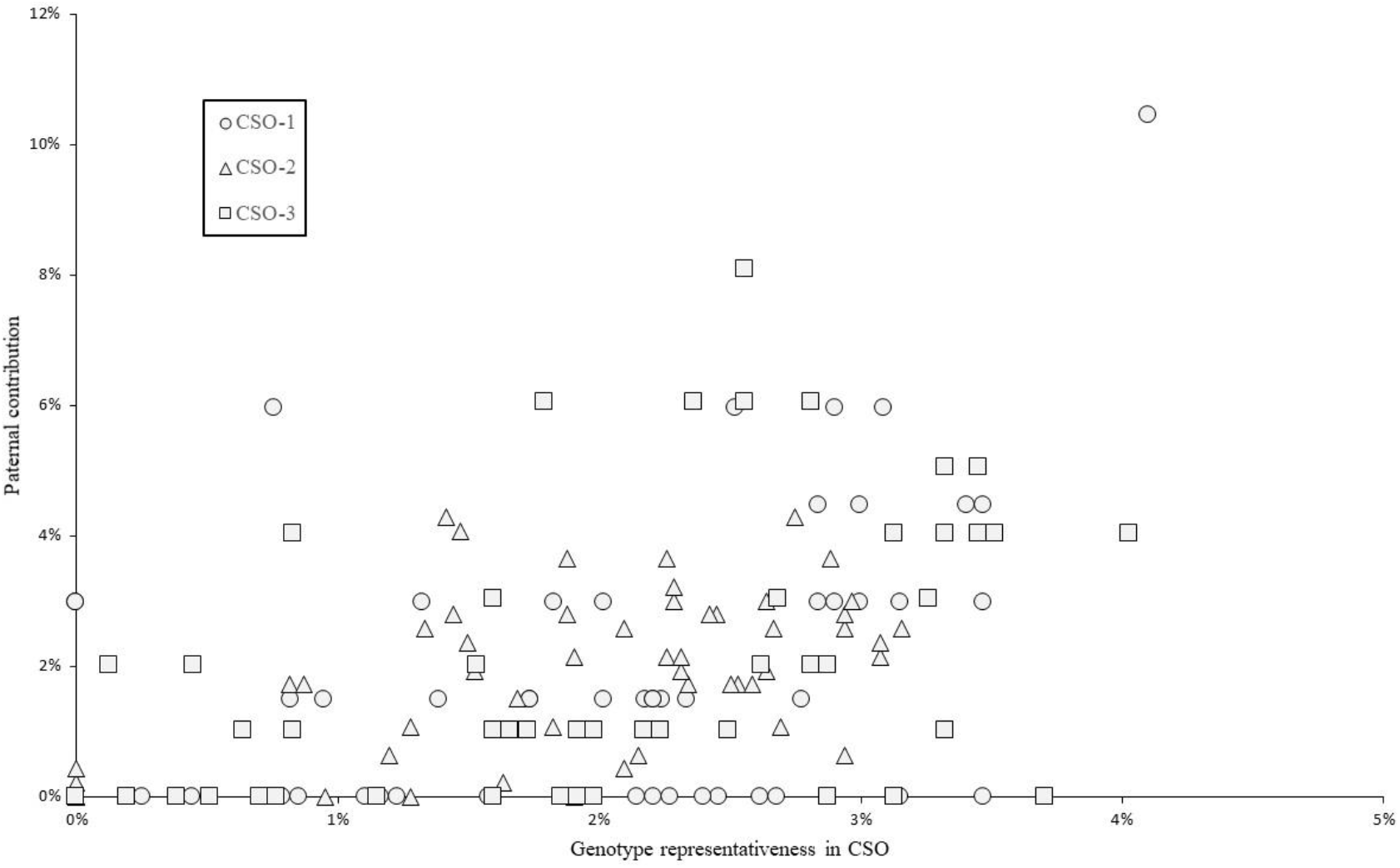
Paternal contribution (% of pollen parents) according to genotype representativeness (% of total number of ramets) in three maritime pine clonal seed orchards (CSO) over 2 pollination years (2011, 2013, sampling strategies SS1 and SS2).

Self-fertilization was estimated for all three sampling sets and amounted to 5.4% over the 2,552 samples analyzed. Results for SS1 are reported in Table 3, with no significant differences detected between the four seed parents (selfing rates were 4.1%, 4.0%, 8.3%, 6.0% for seed parent A, B, C, D, respectively). By contrast, in SS2, selfing rates were variable and ranged from 0 to 26% (Figure 5). The rate of self-fertilization was not correlated with the number of ramets per genotype (data not shown).

**Figure 5.**
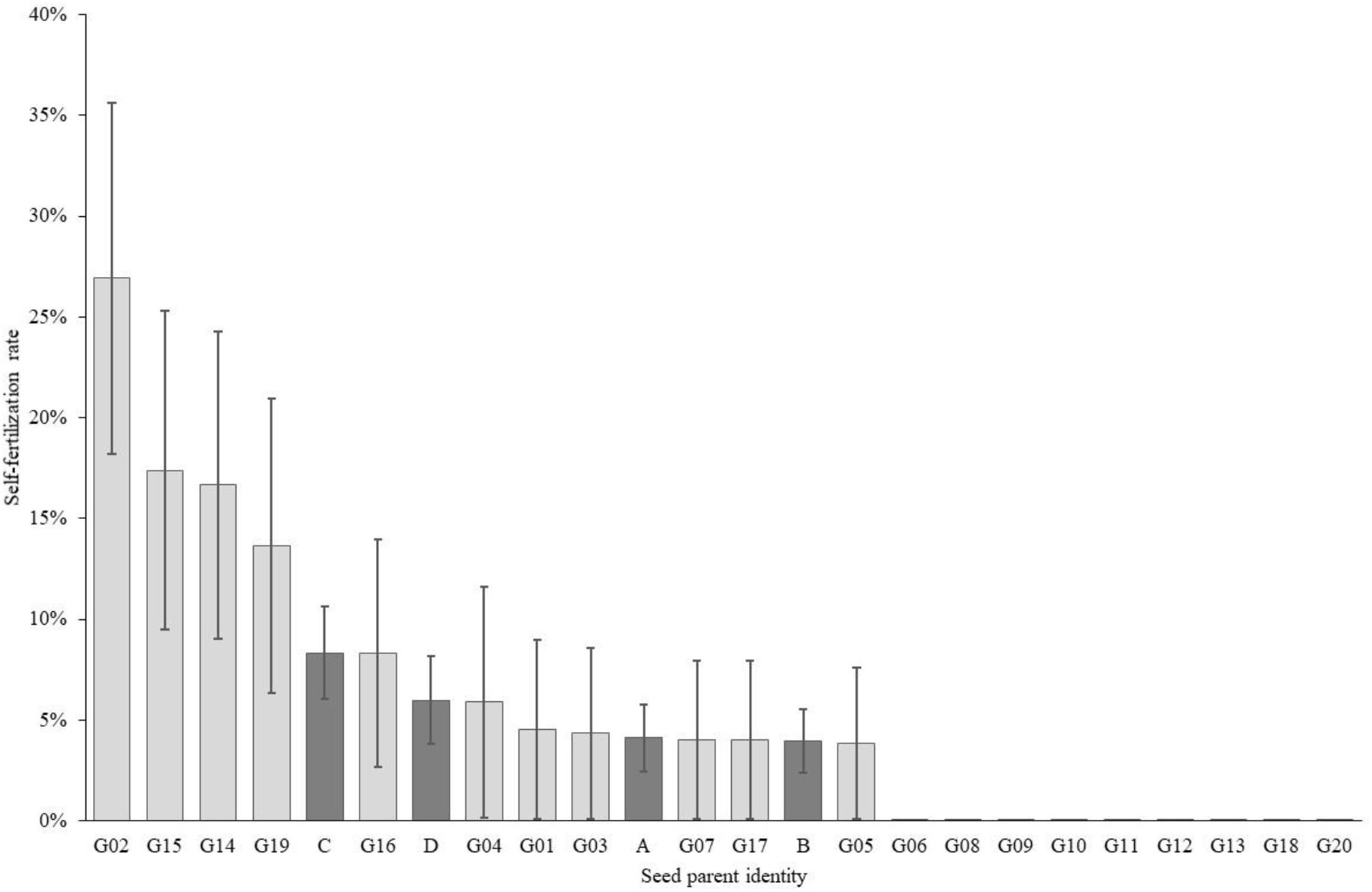
Self-fertilization rates (%) on 24 seed parent genotypes of maritime pine clonal seed orchard CSO-2 pollinated in 2013. Dark grey: four reference seed parents (sampling strategy SS1); Light grey: 20 seed parents (sampling strategy SS2); Bars: standard errors.

### Genetic diversity

Differences between the census number (N) and effective number (N_eff_) of parental genotypes per CSO resulted from the deployment of variable numbers of ramets per genotype (Table 4). Considering only pollen parents from within the CSO, the low N_eff, pollen parent_ (14.6 in CSO-1, 31.2 in CSO-2 and 13.1 in CSO-3) reflected a highly heterogeneous paternal contribution, as shown in Figure 3. The consideration of pollen parents from outside the CSO greatly inflated genetic diversity, particularly in CSO-1 (N_eff, pollen parent_ = 346.9).

**Table 4.**
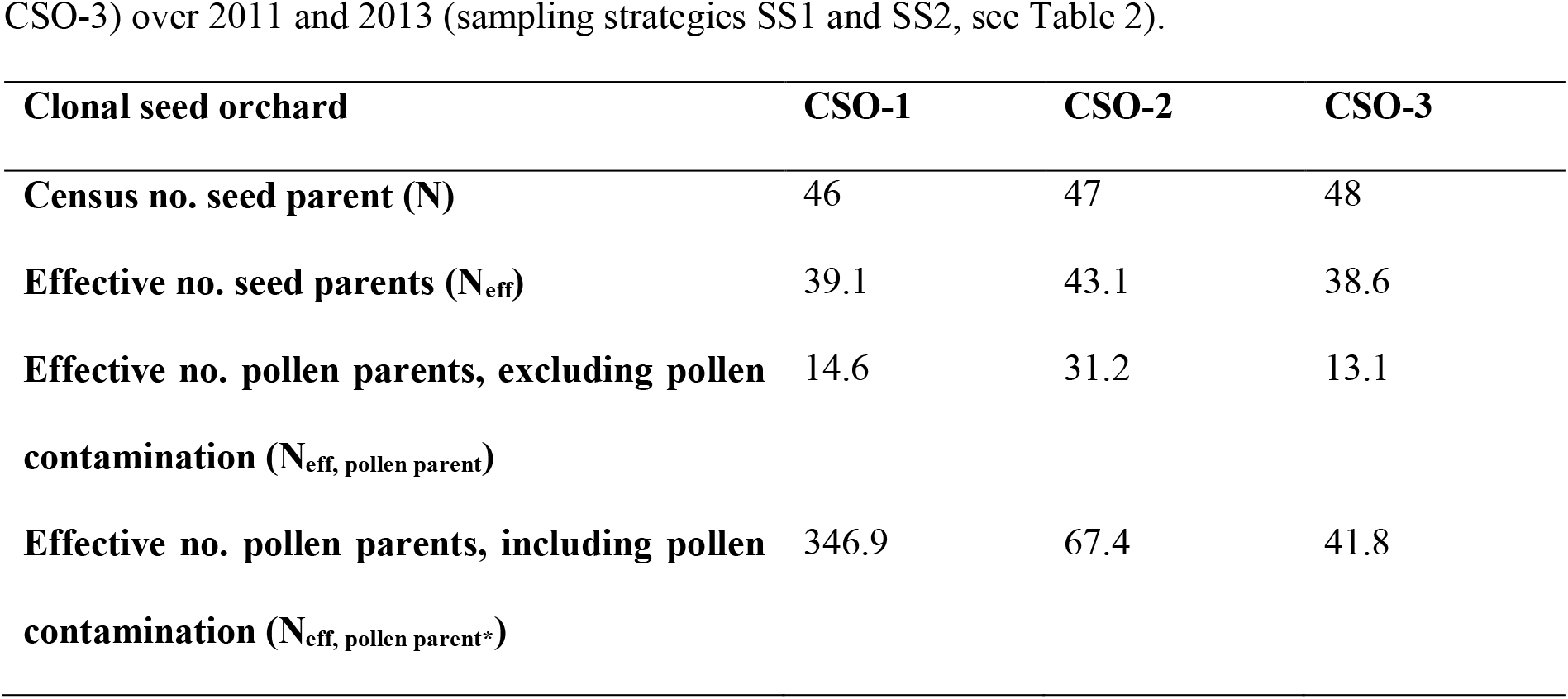
Genetic diversity parameters for three maritime pine clonal seed orchards (CSO-1, CSO-2, CSO-3) over 2011 and 2013 (sampling strategies SS1 and SS2, see Table 2).

| Clonal seed orchard | CSO-1 | CSO-2 | CSO-3 |
| --- | --- | --- | --- |
| **Census no. seed parent (N)** | 46 | 47 | 48 |
| **Effective no. seed parents ($N_{\text{eff}}$)** | 39.1 | 43.1 | 38.6 |
| **Effective no. pollen parents, excluding pollen contamination ($N_{\text{eff, pollen parent}}$)** | 14.6 | 31.2 | 13.1 |
| **Effective no. pollen parents, including pollen contamination ($N_{\text{eff, pollen parent}^*}$)** | 346.9 | 67.4 | 41.8 |

## Discussion

This analysis of 2,552 maritime pine seedlings is, to our knowledge, the largest study to date estimating pollen contamination and mating structure in forest tree seed orchards. Deployment based on open-pollinated CSO is a cost-effective strategy for delivering genetic gains. However, two major assumptions must be satisfied for the objectives of genetic gain and diversity to be fully achieved: i) no contamination with foreign pollen and, ii) random mating between the parental genotypes. Our study reveals a mean pollen contamination rate of 50% and an unbalanced paternal contribution. Based on breeding values estimations (Bouffier et al. 2016), parental genotypes achieve in average 30% genetic gains relative to unimproved material, meaning that their offspring (seed lots) should deliver 30% genetic gains for volume and stem straightness. Assuming that the foreign pollen comes from unimproved stands (thus from trees associated with 0% genetic gains) and an pollen contamination rate of 50%, the expected genetic gain would fall from 30% to 22.5% (50% of the seed lot delivers 30% genetic gains and the remaining 50% which is contaminated with unimproved pollens delivers 15% genetic gains resulting in average to 22.5%), with an accompanying increase in genetic diversity. Unbalanced parental contributions do not induce a systematic bias in genetic gain, but they do decrease genetic diversity. The level of genetic diversity is thus subjected to two adverse forces: pollen contamination and unbalanced parental contribution. However, when high level of pollen contamination is observed, the main driver is the contribution of pollens originated from outside the orchard which inflates the initial level of genetic diversity fixed by the breeder when selecting parental genotypes.

### SNP markers: an efficient tool to study pollen contamination and mating structure

Previous studies (Plomion et al. 2001; Plomion et al. 2005) aimed to estimate pollen contamination in maritime pine seed orchards but cryptic gene flow was suspected due to the low discrimination power of the microsatellite markers available. Moreover, null alleles can be highly detrimental for parentage analyses based on microsatellites (Moriguchi et al. 2004; Plomion et al. 2005; Telfer et al. 2015). Consequently, the reliability of pollen contamination estimated in maritime pine seed orchards with such markers remained questionable. In this study, we used the SNP set developed by Vidal et al. (2015), and demonstrated the power of a set of 60 SNP markers (parental exclusion probability exceeding 99.99%) to estimate pollen contamination rates accurately. Individually, SNP are considered less informative than microsatellites markers, but they are potentially numerous (SNP resources have been published for most conifer species, including maritime pine, Plomion et al. 2016). The main advantages of SNP markers include their high repeatability (Jones et al. 2007), the possibilities for multiplexing and automation of genotyping which makes them cost-effective for high-throughput analyses. In most species, SNP markers have become a tool of choice for parentage analyses (Flanagan and Jones 2019) but only a few studies to date implemented them to estimate pollen contamination in forest trees (Galeano et al. 2021; Hall et al. 2020; Suharyanto et al. 2012).

### High pollen contamination in seed orchards

We found a mean pollen contamination rate of 50%, consistent with the minimum rate of 36% estimated in maritime pine polycross seed orchards by Plomion et al. (2001), using chloroplast microsatellites. Plomion et al. (2005) subsequently used nuclear microsatellites and revealed similar high pollen contamination rates in second-generation seed orchards (32% to 81%). Medium-to-high levels of pollen contamination have been reported in conifer species: 28% in *Pinus thumbergii* (Suharyanto et al. 2012), 49% in *Cryptomeria japonica* (Moriguchi et al. 2010), 12-35% in *Pseudotsuga menziesi* (Korecký and El-Kassaby 2016; Slavov et al. 2005), 27% in *Picea glauca* (Galeano et al. 2021), 58% in *Picea abies* (Dering et al. 2014), 5-52% in *Pinus sylvestris* (Funda et al. 2015; Torimaru et al. 2009), and 86% in *Pinus brutia* (Kaya et al. 2006). The various seed lots and sampling strategies as well as the large number of seedlings analyzed in this study allow us to explore various biological and environmental factors to explain such high pollen contamination rates in seed orchards.

First, the location of the CSO emerged, as expected, as a key factor explaining pollen contamination due to the vicinity with the source of external maritime pine pollen (unimproved stands). *In situ* experimental studies in a maritime pine seed orchard showed that about 20% of pollen contamination could be explained by distant pollen flows, the remaining 80% being explained by local input within a range of ten to several hundred meters (Baradat et al. 1984; Castaing and Vergeron 1976). In other conifer species, such as Scots pine and loblolly pine, viable pollen can travel dozens or even hundreds of kilometers (reviewed by Kremer et al. 2012). Assuming a similar pollen dispersal profile in maritime pine, CSO-1 (located within the forest) would experience massive local pollen flow from the Landes de Gascogne forest, whereas CSO-2 (outskirts of the forest) and CSO-3 (outside the forest) would potentially receive more limited pollen flow from this source. Interestingly, similar contamination rates were detected in 2011 and 2013 in the center and at the edge of CSO-1 confirming the results published by Funda et al. (2015) in Scots pine and suggesting that the whole orchard is subject to homogeneous contamination with outside pollen due to long-distance pollen flows. In addition, the soil conditions of the three seed orchards are rather different (sandy soil for CSO-1 vs clay loam soil for CSO-2 and CSO-3). Clay loam soils are known in maritime pine to be associated with the earlier formation of strobili, about 7-10 days ahead of most of the Landes de Gascogne forest located on sandy soils. The receptivity of the female strobili in CSO-2 and CSO-3 may therefore be optimal well before the emission of massive amounts of pollen from the Landes de Gascogne forest, in which CSO-1 is located. However, both effects (vicinity with external pollen sources and soil conditions) are cofounded in our study which makes it impossible to estimate the relative importance of each of them.

Second, the intensity of flowering increases with tree age and becomes optimal for commercial harvesting after about 8-10 years. CSO-1 has been planted 3-4 years earlier (2006) than CSO-2 and CSO-3 (2002-2003). At the time of first sampling in our experiments (2011), a lower rate of fertile male and female strobilus is therefore likely in CSO-1 (5 years old) compared to CSO-2 and CSO-3 (8-9 years old). Internal pollen flows at CSO-1 could be insufficient to compete with massive external sources and result in very high rates of pollen contamination (96% in 2011). Accordingly, pollen contamination observed two and three years later in CSO-1 was reduced (60% and 70%). However, contamination levels can remain high in old seed orchards, as highlighted by Torimaru et al. (2009). This is consistent with the rates observed in CSO-2 and CSO-3 which remain at quite high level in 2013 and 2014 (20-35%, age 10-12).

Third, beyond the tree maturity discussed above, meteorological factors could be involved in the annual variability of pollen contamination which is higher in 2011 compared to 2013 and 2014, whatever the CSO considered. Temperature, rainfall, wind strength and direction during flowering could affect the formation, persistence, and outcome of pollen clouds, as well as the viability of pollen released in spring. Mean daily rainfall during the period of female strobilus receptivity (estimated from control crosses in the framework of the maritime pine breeding program) was 0.5 mm in 2011 vs. 1.9 mm in 2013 and 2.4 mm in 2014 (Météo-France data). Dry periods, such as that observed in 2011, favor pollen flow over long distances, as rain is known to affect the extent of pollen dispersal (Di-Giovanni and Kevan 1991).

Finally, the identity of the seed parent could affect the flowering. We do observe a high variability in pollen contamination rates depending on genotype (Figure 2). However, unlike Slavov et al. (2005), who reported higher contamination rates for genotypes with early female receptivity, we found no relationship between the timing of pollen receptivity (estimated during pollination years 2015 and 2016) and pollen contamination (Trontin et al. 2019). Our findings suggest that the female flowering phenology within the orchard had little impact on pollen contamination, probably due to the extended period of pollen release. The four seed parents sampled for SS1 were ranked among the most contaminated seed parents sampled in SS2 (Figure 2). We currently have no explanation for this observation, but it may have biased the pollen contamination rates estimated in 2011 and 2013 upwards in the three CSO.

### Uneven paternal contribution to seed lots and variable self-fertilization rate

The sampling strategy used here, based on the collection of cones from specific seed parents, were not designed to study the contribution of the seed parents, but it was possible to analyze paternal contribution based on SS1 and SS2. All pollen parents were recovered at least once in the seed lots genotyped, but a high level of variability was observed for paternal contribution, as also reported by Suharyanto et al. (2012) in *Pinus thunbergii.* The weak correlation between genotype representativeness (based on the number of ramets per genotype) and paternal contribution (Figure 3) suggests that genotypes released various amounts of pollen, as confirmed by Trontin et al. (2019). Seed orchard design is optimized to minimize self-fertilization (which leads to inbreeding depression in conifer species) by spatially separating copies of the same genotype. The overall rate of self-fertilization was estimated at 5.4% at the seedling stage, a value below the 13% reported by Baradat et al. (1984) for maritime pine, but within the range of estimates for pine seed orchards (Funda et al. 2015; Suharyanto et al. 2012; Torimaru et al. 2009). As previously reported by Funda et al. (2015), self-fertilization rates depended strongly on seed parent identity and was as high as 26.9% for one genotype in our study.

The uneven parental contribution in the seed lots tends to decrease genetic diversity (Table 4) but, when considering the effect of pollen contamination, the final effective number of parents is generally higher than the one initially set by the breeder.

### Towards optimized deployment of maritime pine seed orchards

Various management practices have been proposed for reducing pollen contamination in forest tree seed orchards. These practices include supplemental mass pollination (Korecký and El-Kassaby 2016; Stoehr et al. 2006), water cooling to delay strobilus production (El-Kassaby and Davidson 1991; Song et al. 2018) and greenhouse-like structures (Funda et al. 2016; Moriguchi et al. 2010; Torimaru et al. 2013). Our study reveals that pollen contamination in French maritime pine CSO could be strongly reduced by:

i) choosing the location of the orchard carefully, in terms of its distance from external pollen sources and soil conditions, ii) not collecting seeds from young trees (below 8 years old). The methodology used here, based on a set of 60 SNP markers, proved cost-effective and highly powerful for parentage reconstruction. Our results suggest that sampling 100 seeds annually should be sufficient to estimate pollen contamination (this sample size provides estimates with a standard error of 5%) for both applied uses (seed lot quality certification) and for research purposes (e.g. exploring links between pollen contamination and climatic data: yearly variations in pollen contamination may be associated with rainfall levels during the pollination period). Finally, flowering phenology, as well as pollen and cone productivity are known to be under strong genetic control in conifers (Wu et al. 2021). A better knowledge of these flowering traits in the whole breeding population is required, to optimize seed orchard composition and to hone estimates of the expected genetic gain.

## Author contributions

LB and JFT conceived and designed the study with expert support of PA, AR and PP. LB coordinated the study. PA collected samples. SD contributed to the molecular laboratory work. LB carried out the genetic analyses and drafted the first version of the manuscript with contributions from SD and JFT. All authors contributed to the discussion and approved the final version of the manuscript.

## Data availability

Information about seedlings (CSO, pollination year, sampling strategy, sampling zone, seed parent genotype), SNP description, as well as genotyping data for the 2,552 seedlings and the 51 CSO parents are available from https://doi.org/10.57745/SR2HAJ

## Acknowledgments

This study was supported by a national grant (QUASEGRAINE project, French Ministry of Agriculture/DGAL, no. 2014-352, coordinated by ONF/B. Musch) and regional funds from the Conseil Régional d’Aquitaine (IMAF project, no. 12009468-052, coordinated by FCBA/L. Harvengt) and the Conseil Régional Centre Val de Loire (IMTEMPERIES project, no. 2014-00094511, coordinated by INRAE/M.-A. Lelu-Walter). We thank Vilmorin and Forelite for providing access to the orchards and the Maritime Pine Breeding Cooperative (GIS Groupe Pin Maritime du Futur) for its support through the FORTIUS project (grants from the Conseil Régional d’Aquitaine and the French Ministry of Agriculture, coordinated by INRAE/P. Pastuszka). Samples were prepared at Xylobiotech (https://www6.inrae.fr/in-sylva-france/Services/In-Lab/XYLOBIOTECH) — the shared FCBA/INRAE platform dedicated to forest biotechnology supported by the ANR (ANR-10-EQPX-16 Xyloforest) — with contributions by Francis Canlet (FCBA Research Technician), Marie Chambard and Mathilde Staat (FCBA trainees). The SNP genotyping was performed at the Bordeaux Genome Transcriptome Facility (doi:10.15454/1.5572396583599417E12).

The authors would like to dedicate a special tribute to Marjorie Vidal who passed away in 2021. Marjorie’s PhD work (2016, supervision INRAE/FCBA, University of Bordeaux) has notably contributed to the development of SNP arrays for pedigree reconstruction in maritime pine (Vidal et al. 2015).

## Abbreviations

CSO: clonal seed orchard (three CSO were analyzed in this study: CSO-1, CSO-2, CSO-3)
FRM: Forest Reproductive Material
RAPD: Random Amplified Polymorphic DNA
SNP: single-nucleotide polymorphism
SS: sampling strategy (three SS were considered in this study: SS1, SS2, SS3) A, B, C, D: name of the four reference seed parents selected for SS1 based on contrasted flowering phenology

